# Genetic Structure and Diversity of Amaranth Populations across Multi-Environmental Sites in Burkina Faso Revealed by Simple Sequence Repeats (SSRs) and Genotyping-by-Sequencing (GBS)

**DOI:** 10.64898/2026.01.09.697177

**Authors:** T. Tenhola-Roininen, O. Bitz, L. Bitz, S. Kankaanpää, S. Rokka, Z. Kiebre, K.R. Nanema, H. El Bilali, V.-M. Rokka

## Abstract

The genus *Amaranthus* L. (Amaranthaceae) comprises 70–80 species, including leafy vegetables, pseudocereals, ornamentals, and weedy forms, and is widely distributed across tropical and temperate regions. Despite its nutritional and agronomic importance, the taxonomy of *Amaranthus* remains challenging due to extensive phenotypic plasticity, sporadic hybridization, and limited diagnostic traits. In Burkina Faso, three species (*A. cruentus, A. hypochondriacus*, and *A. dubius*) are commonly cultivated, displaying considerable morphological variability and forming seven distinct morphotypes. To better understand the genetic diversity and population structure of West African *Amaranthus*, we analyzed accessions from multi-environmental sites in Burkina Faso using both simple sequence repeat (SSR) markers and genotyping-by-sequencing (GBS). SSR-based STRUCTURE analysis revealed a pronounced ΔK peak at K = 2, indicating two major genetic groups, with additional substructure at K = 4. Principal Coordinates Analysis confirmed species-level separation and morphotype-based clustering, while bar plots from STRUCTURE analysis highlighted admixture among several accessions. Complementary GBS analysis provided higher-resolution insights, partitioning species into multiple sub-clusters, detecting introgression, and linking genetic differentiation to morphological traits such as leaf shape, pigmentation, and inflorescence type. These findings are consistent with previous studies in Burkina Faso that reported both major genetic groups and significant morphological diversity. Together, SSR and GBS analyses underscore the multilayered nature of *Amaranthus* diversity, with SSRs capturing broad population subdivision and GBS refining fine-scale structure. This dual approach provides a robust framework for conservation of genetic resources and for genomics-assisted breeding aimed at exploiting heterosis, preserving unique alleles, and developing resilient varieties to enhance food and nutrition security in sub-Saharan Africa.

## Introduction

The amaranth family (*Amaranthaceae* Juss.), includes about 165 genera and over 2,000 herbaceous plants, making it the most species-rich group within the order *Caryophyllales. Amaranthus* L. is a genus of comprising 70-80 species (Iamonico et al. 2023), and it includes mostly wild and weedy, but also some domesticated vegetable, grain crop and ornamental species (Costea et al. 2001). This genus is widely distributed across tropical and temperate regions of the Americas, Africa, Asia, and Europe. Suma et al. (2002) reported that from those about 70 *Amaranthus* species, 400 are varieties distributed throughout the world with only a few being domesticated in different countries. The species of *Amaranthus* are often difficult to characterize because of their similarities and existence of intermediate forms (Aderibigbe et al. 2022). Their taxonomic differentiation is difficult because only few traits are suitable for this purpose despite a high phenotypic diversity (Stetter & Schmid 2017). In addition, the taxonomy of *Amaranthus* is not well understood due to its wide phenotypic variation, which is partly caused by sporadic hybridization between different species. A small number of suitable traits, phenotypic plasticity, gene flow and hybridization have made it difficult to establish the taxonomy and phylogeny of the whole *Amaranthus* genus despite various studies already made using molecular markers (Stetter and Schmid 2017).

The *Amaranthus* species are mostly selfing, diploids (2n=2x=32, 2n=2x=34) with genome size 500 Mbp (Stetter and Schmid 2017). *Amaranthus dubius* may also exist as tetraploid (2n=4x=64) being the only native tetraploid amaranth species known so far https://prota.prota4u.org. Grain amaranths are originated from Central and South America, where they were of great importance in local agriculture until their cultivation strongly declined after the Spanish conquest (Stetter et al. 2016). Mainly three species of *Amaranthus* are cultivated as pseudocereals for grain production: *A. caudatus* L., *A. cruentus* L., and *A. hypochondriacus* L. (Stetter et al. 2016). In many tropical and subtropical regions, edible leaf-amaranths are also a significant source of protein and micronutrients and often substitute for or complement spinach-like vegetables. About 20–30 species of *Amaranthus* are used as leafy vegetables worldwide, and especially *A. blitum, A. tricolor, A. cruentus, A. dubius, A. edulis*, and *A. hypochondriacus* are the species whose leaves are used for nutrition (Andini et al. 2018). In some African societies, protein from *Amaranth* leaves provides even 25% of the daily protein intake during the harvest season (National Research Council 2006).

The global food system has been dependent on only a few plant and animal species and strains (Khoury et al. 2014), and there are considerable challenges in achieving food and nutrition security in diverse regions (Sarker et al. 2022). Therefore, also NUS (neglected and underutilized species) have an essential role for maintenance of global food security and are extremely important for food production especially in Low Income Food Deficient Countries (LIFDCs) (Das 2012). NUS offer new opportunities to address food insecurity and malnutrition that are aggravated by population growth, climate change and land degradation (Bilali 2020). Amaranth is one of such crops with high genetic diversity and it is well adapted to drought and marginal land (Alemayehu et al., 2015). Amaranths adapt very well in changing osmotic conditions as *Amaranthus* species use C4 pathway for photosynthesis using CO_2_ efficiently in different ranges of temperature and water stress conditions (Liu and Stützel 2002). It is a promising vegetable crop species grown under semi-arid conditions, where the nutritional benefits and drought resistance are critical. The high adaptations of amaranths may be a reason for its geographic spreading in many parts of the world (Khamis et al. 2025). Amaranth demonstrates resistance to drought, heat, and pests, and adapts more readily to harsh environments than conventional cereals. These traits contribute to its growing reputation as a crop of even high economic importance for challenging climates (Yadav & Yadav 2024).

In Burkina Faso, Western Africa, amaranths are locally known as “Bolombourou” by the local Mooré language. Six *Amaranthus* species have been identified in the region (*A. dubius, A. graecizans, A. cruentus, A. hypochondriacus, A. viridis*, and *A. spinosus*) (Ouedraogo et al. 2011), but according to Ouedraogo et al. (2021) only three species are commonly cultivated in Burkina Faso: *A. cruentus, A. hypochondriacus*, and *A. dubius*. The cultivated Amaranth species are generally divided into two groups; those grown for their edible leaves and those grown for their seeds (Ouedraogo et al. 2021). Among the grain types, *A. hypochondriacus* stands out as the most robust and productive, well-suited to tropical and dry climates. Previous studies in Burkina Faso have shown great morphological variability within cultivated species and have identified seven morphotypes whose main classification criteria have been the color and shape of leaves and inflorescences (Ouedraogo et al. 2021, 2024). The green-leaved *A. cruentus* has the highest leaf biomass yield, a long flowering cycle, and makes a high number of branches. Early-flowering, oval-leaved accessions with broad, long blades have been found in *A. dubius* and this species is grown exclusively for its leafy greens in Burkina Faso (Ouedraogo et al. 2021). In addition, *A. cruentus* serves dual purposes, as a pseudocereal and a leafy vegetable. The green morphotype of *A. cruentus*, particularly the variant with black grains, is the most widely cultivated and consumed amaranth in Burkina Faso and in neighboring West African countries such as Benin, Ivory Coast, Togo, Nigeria, and Senegal (Aderibigbe et al. 2022).

Genetic variation holds the key to the ability of populations and species to persist over evolutionary period through changing environments. Those plant populations that have a narrow range of genotypes and are more uniform may merely fail to survive and reproduce at all, when the conditions become locally less favorable. In small out-cross populations, severe reductions in population sizes cause loss of genetic diversity and increased susceptibility to infectious pests and diseases. Keeping the genetic resources for cultivated and crop species, such as in NUS, plays an important role in providing adaptive and productive genes, thus leading to long-term increases in food productivity. Genetic diversity analyses in *Amaranthus* are essential for understanding its evolutionary relationships, domestication history, and potential for crop improvement. Assessing genetic variation across wild species, landraces and cultivars helps researchers trace domestication pathways, identify ancient centers of origin, and clarify how human selection shaped agronomic traits such as seed size, nutritional composition, and stress tolerance.

Microsatellite (SSR) marker analysis and genotyping-by-sequencing (GBS) each offer distinct advantages for assessing genetic variation, population structure, and overall diversity. Microsatellites provide highly polymorphic, co-dominant markers that enable accurate discrimination among genotypes and are cost-effective for studies with moderate sample sizes. Their established laboratory protocols and cross-study comparability further enhance their utility in population genetic analyses. In contrast, GBS offers genome-wide, high-throughput SNP discovery without the need for prior genomic resources, producing large marker datasets that enable fine-scale resolution of genetic diversity, rare variant detection, and detailed inference of population structure and evolutionary relationships. Together, these approaches provide complementary strengths for comprehensive genetic characterization.

In the present study, the genetic structures of Western-African *Amaranthus* seed collections originated from multi-environmental sites of Burkina Faso were analyzed. The materials included previously identified/non-identified *Amaranth* populations with at least seven morphotypes, and both SSR marker analysis and GBS method were applied.

## Materials and Methods

### Plant material

A total number of 91 accessions from three different *Amaranthus* species (*A. cruentus, A. hypochondriacus, A. dupius*) according to the taxonomic identifications made by Ouedraogo et al. (2011, 2021, and 2024) were delivered from UJKZ (Université Joseph KI-ZERBO), Ouagadougou, Burkina Faso. The seeds for amaranth accessions were collected from growers at 11 market garden sites in different climatic zones of Burkina Faso. A total of 19 accessions were collected in three provinces (Bam, Séno, Yatenga) of the Sahelian climatic zone (semi-arid dry region), 47 accessions collected in seven provinces (Bazèga, Boulkiendé, Gourma, Kadiogo, Kouritenga, Oubritenga, Sanguié) of the Sudano-Sahelian zone (transitional region), and 17 accessions collected in the province of Houet located in the Sudanese zone (wetter region) **(Figure 1.)**. In addition, 7 accessions (Wolveg) included into the study were derived from World Vegetable Center. The annual rainfalls in Sahelian zone of Burkina Faso are 300-600 mm, in Sudano-Sahelian zone 600-900 mm, and in Sudanese zone 900-1,200 mm.

**Figure 1.**
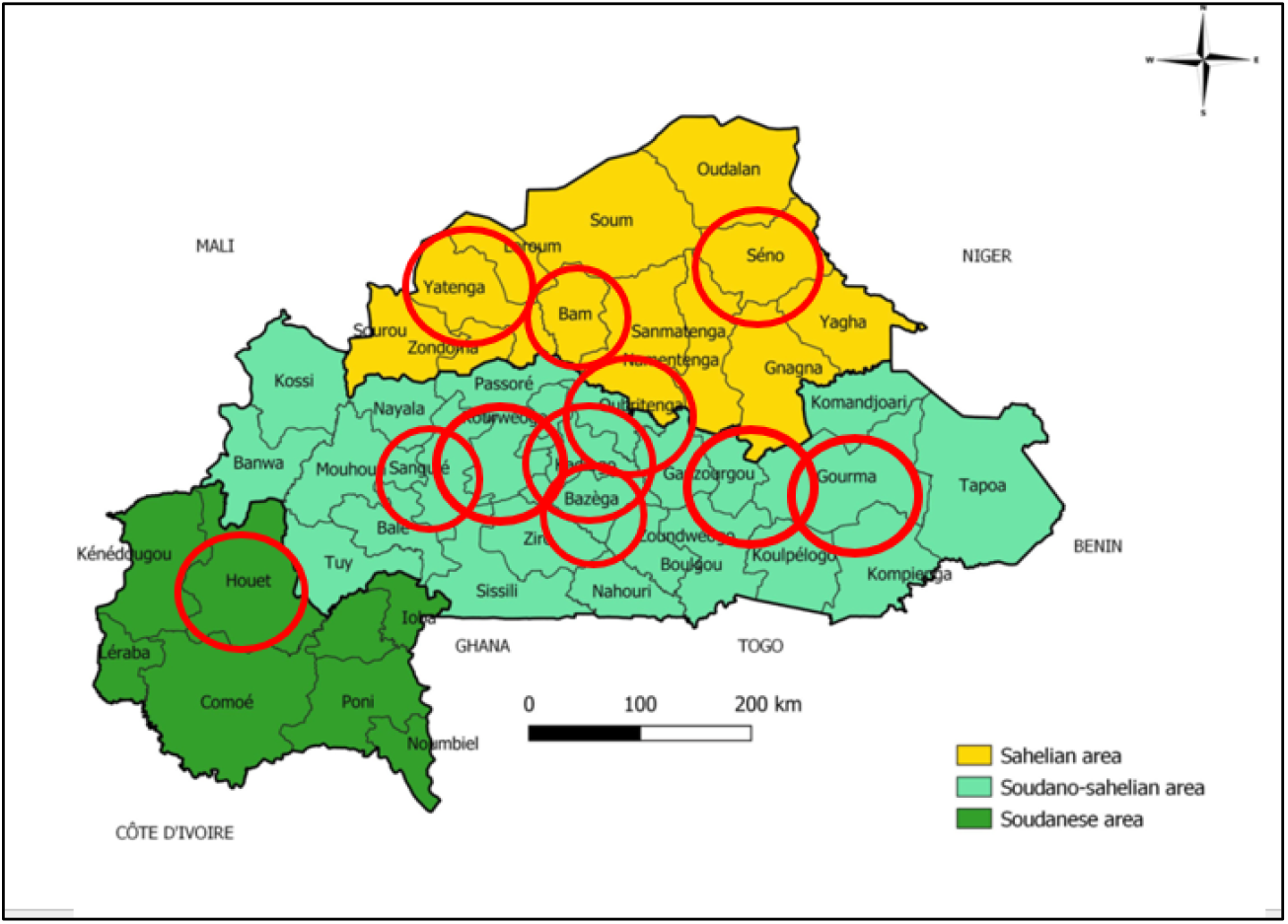
The eleven provinces of Burkina Faso, where the *Amaranthus* accessions were collected, marked by red circles. Three provinces (Bam, Séno, Yatenga) are in the Sahelian zone (yellow), seven provinces (Bazèga, Boulkiendé, Gourma, Kadiogo, Kouritenga, Oubritenga, Sanguié) in the Sudano-Sahelian zone, and the province of Houet located in the Sudanese zone.

In this study amaranth populations consist of six morphotypes (Table 1.). They were classified as light green (3 accessions of *A. hypochondriacus*), green (31 accessions of *A. cruentus*), dark green (3 accessions of *A. dubius*, 4 accessions of *A. hypochondriacus*, 1 accession from Wolveg collection), green purple (1 accessions of *A. hypochondriacus*), green violet (1 accession of *A. cruentus*), and violet (5 accessions of *A. cruentus*). In total 14 accessions were included to the study without knowledge of morphotype/species identification. The seeds of amaranth accessions were delivered from Burkina Faso to Finland in May 2023. Seeds were sown in the growth chamber with temperature range of 23-25 °C/18 °C (day/night), 16 h daylength, and grown until five-leaf stage. From all accessions germinated, leaves were collected and frozen at – 20 °C. For genetic diversity analyses, 60 different amaranth accessions from 91 accessions obtained from Burkina Faso were studied **(Table 1)**. In every population, three to six specimens were analysed in the microsatellite and only one in genotyping by sequencing (GBS) studies.

Table 1. As attachment

### DNA extractions

Before DNA isolation, the frozen leaves (30-50 mg) of amaranth samples were homogenized using Tissuelyser II (Qiagen, USA) for 1/25 m/s speed for 2x 45 s with 6mm steel beads including 60 s cool down period in between, and then extracted with Qiacube HT 96, automated isolation system, using DNeasy Plant Kit (Qiagen, USA) according to manufacturer’s protocol. In some cases, vacuum was not successful eluting all liquid and then excess was discarded before allowing the protocol to proceed into the next step. DNA was eluted into 100 µl of milli-Q water and quantified using 8-channel Nanodrop (Thermo Fischer Scientific). Across the analyzed samples, the DNA concentrations varied, the mean value being 31 ng/µl. Three to six specimens per accession were used in microsatellite analysis. Single samples from each accession could be used for GBS-library preparation since supportive data was already available from microsatellite data (Supplemental Table 1).

### Microsatellite analysis

Eight polymorphic SSR primers (Table 2.) were chosen for the diversity analyses (Tiwari et al. 2021; Lee et al. 2008). Polymerase chain reaction (PCR) program was modified according to protocols by Tiwari et al. (2021) and Lee et al. (2008), respectively. The total volume of PCR mixture (10 µl) contained 10 ng of DNA (2 ul), 5 pmol (1 µl) primer pairs, 1U (0.2 µl) DNA polymerase (Biotools, B & M Labs, S.A., Madrid, Spain, 5U/µl), 1x buffer (1 µl) (Biotools B & M Labs, S.A., Madrid, Spain, 10x) including 2mM MgCl2, and 200 µM (1 µl 2 mM) dNTPs. In PCR mixtures, primer pairs were pooled to form multiplexes 1 and 2 (Table 2). The tail of each forward primer was labeled with fluorescent dye, FAM, VIC, or NED (Applied Biosystems UK, Life Technologies Europe BV (Table 2). In all PCR reactions, annealing temperature was 55°C. The PCR reaction started with denaturation for 4 min at 94 °C, then cycles 35 times; denaturation 30 s at 94 °C, annealing 30 s at 55 °C and last cycle step 60 s at 72 °C, and final extension step 10 min at 72 °C before cooling down to 4°C. All PCR products with two multiplexes were analyzed using the ABI 3500XL Genetic analyzer (Thermo Fischer Scientific Inc., Vantaa, Finland) with Sanger sequencing system. The PCR products were diluted 1/20- 1/50 before the ABI runs.

**Table 2.**
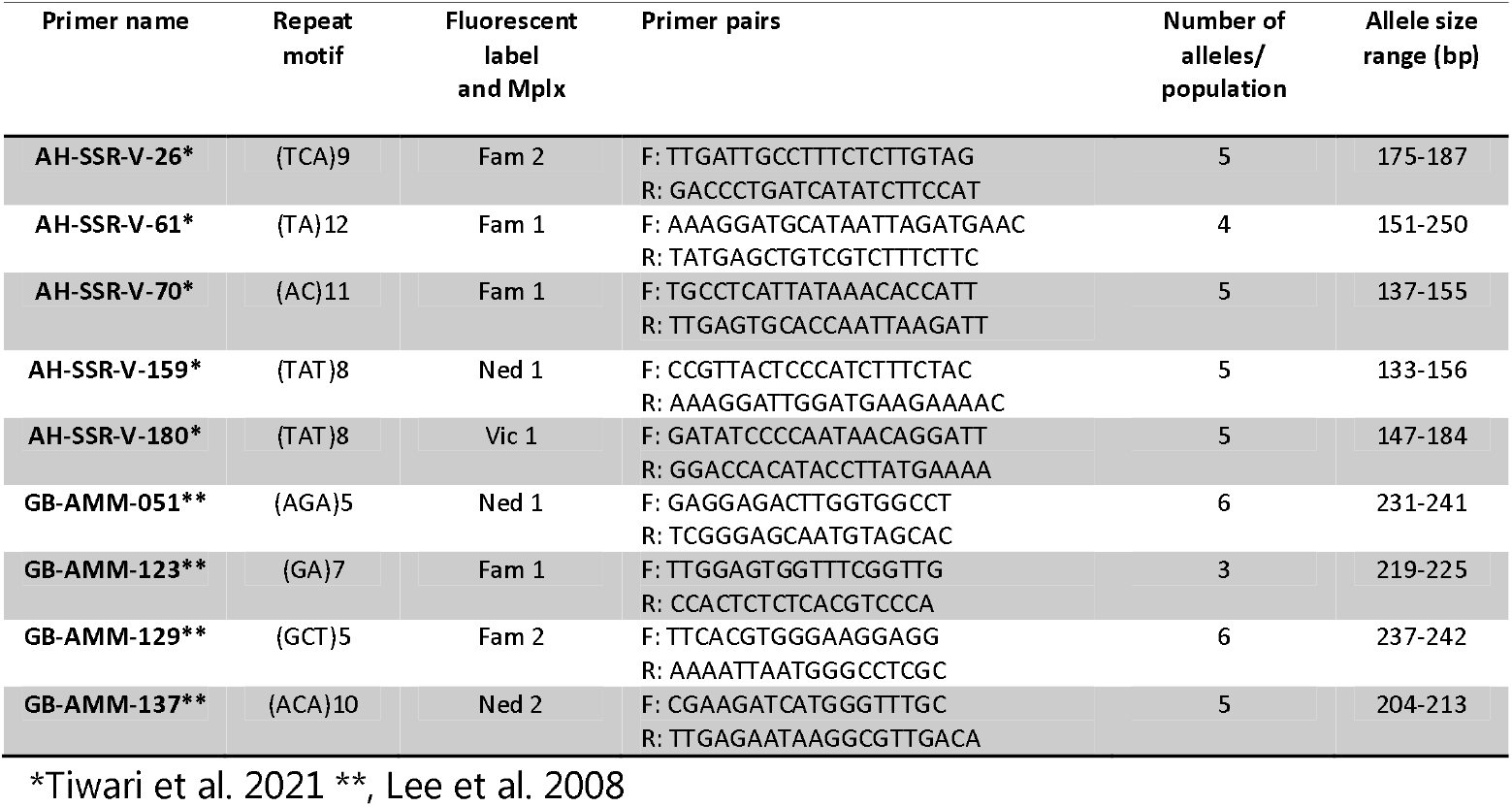
SSR primer pairs used in the diversity study.

### Diversity analyses for the microsatellite data…

GeneMapper version 6 (Applied Biosystems) was used for allele analyses. Genetic distances between samples were analysed by using un-weightened neighbor joining method in DarWin version 6.018. Samples with a lot of missing data >30% were excluded from the dendrogram analyses. The program GenAlex version 6.5 was used for principal component analyses (PCoA), in which Nei’s genetic distances were used for analyses. Analysis of molecular variance (AMOVA) was performed using GenALex 6.5 (RESULTS MISSING) Structure version 2.3.4 (Pritchard et al. 2000) was used for Bayesian clustering analyses. The analysis was run for values of K ranging from 1 to 10, with 10 independent replicates per K. Each run employed a burn-in period of 150,000 iterations followed by 200,000 MCMC repetitions under the admixture model with correlated allele frequencies. In the run, populations were not predefined. Structure Selector (2018) was used for calculating the best delta K value and creation coefficient matrices (Q-matrices) by CLUMPAK (2015) which identifies clustering modes and analyses population structure across K. Distruct version 1.1. generated the visual interpretation.

### ddRAD library preparation

58 specimens from 58 different amaranth populations and several locations were used for GBS analysis. The optimized protocol (Fischer et al. 2025) for ddRAD library preparation was adapted from Salas-Lizana & Oono (2018). Briefly, high quality DNA was double-digested with two restriction enzymes PstI-HF and MspI (NEB, Bionordika, Finland). The restriction reaction was performed in a volume of 20 µL, containing 300 ng of DNA, 0.25 µL of PstI-HF (5 units), 0.5 µL of MspI-HF (10 units), 2 µL of cut-smart buffer (NEB, Bionordika, Finland) (10x) and 0.25 µL of molecular grade water at 37°C for 3 h, following heat-inactivation for 15 min at 65°C. The subsequent ligation of two non-barcoded restriction site specific adapters (Table) was performed by adding to the restriction mixture 1 µL of each adapter, PstI 1 µM and MspI 10 µM, 0.4 µL of T4 ligase (NEB, Bionordika, Finland) and 1.5 µL of ligation buffer (NEB, Bionordika, Finland). Ligation was performed at 16°C for 14 h, following heat-inactivation at 65°C for 15 min. DNA-restriction fragments were selected between 175 bp and 700 bp by using SPRIselect magnetic beads (Beckman Coulter, USA) with a left-right ratio of 0.9x-0.56x. The volume of each sample was adjusted with molecular grade water to 50 µL and then 28 µL of SPRIselect beads were added to achieve a 0.56x ratio for the selection of fragments shorter than 700 bp following selection of fragments longer than 200 bp by adding 22 µL of SPRIselect beads to achieve a ratio of 1x. The size selected DNA was resuspended in 25 µL of molecular grade water. Samples were barcoded by adding Illumina Nextera v2 (Illumina, San Diego, CA, USA) combinatorial dual-indexed barcodes (i7 and i5). For each individual sample, the following PCR-mix was prepared: 6 µL of 5x Phusion HF buffer, 0.4 µL dNTP (10 mM), 0.2 µL of Phusion HF DNA polymerase (0.4 units) (ThermoFisher scientific, USA), 1.5 µL of i5 barcode primer, 1.5 µL of i7 barcode primer, 5 µL of sample and 15.4 µL of molecular grade water. Two PCR reactions per sample were performed. The cycling conditions were as follows: initial denaturation at 98°C for 30 sec, followed by 18 cycles of 10 sec at 98°C, 20 sec at 61°C, 15 sec at 72°C and a final elongation step at 72°C for 10 min. The two PCR reactions per sample were pooled and quantified using Qubit Flex with 1x dsDNA HS assay (ThermoFisher scientific, USA). Only products with a significantly higher amount than the No Template Control (NTC) were used for sequencing (>3 ng/µL).

### Sequencing

Single ddRAD libraries were pooled in equimolar amounts, cleaned with 1 x SPRi select beads and resuspended in 30 µL molecular grade water. The pool was selected utilizing BluePippin automated electrophoresis system (Sage Science, distributed by BioNordika Finland Oy; Finland) to the length between 300 and 600 bp corresponding to the combined length of 150-450 (550) bp restriction insert and 147 bp adapter. Following cleanup with 1x SPRiselect beads the library were resuspended in 25 µL molecular grade water. The quality and size of the pooled sequencing library was evaluated on the TapeStation 4150 (Agilent, USA) using the DNA HS1000 assay. Quantification of the library was done using Qubit 4 (1x dsDNA HS assay) (ThermoFisher scientific, USA). Sequencing was done at the institute of Biotechnology/University of Helsinki (Bidgen) on the AVITI24 (Element bioscience) in Helsinki.

### Bioinformatics

After demultiplexing, FastQC was run on each sample and summarized with MultiQC. Only samples with more than 100.000 cluster were kept for the subsequent variant calling (VCF) utilizing the in-house developed Snakemake pipeline Snakebite-GB (Fischer et al. 2025), available on GitHub (https://github.com/fischuu). Minor allele frequency was set to 0.01. *Amaranthus cruentus* (GCA_048987545.1_ASM4898754v1) was used as reference genome. The samples used to build the mock reference were randomly selected samples from each species of Amaranth: KOMB2 (*A. cruentus*), KAD4 (*A. dubius*) and SAN13 (*A. hypochondriacus or cruentus*). fastreeR (https://www.bioconductor.org/packages/release/bioc/html/fastreeR.html) as used to calculate the distances and Archaeopteryx ([http://www.phylosoft.org/archaeopteryx) to visualize the dendrogram.

## Results

To investigate the genetic structure of the *Amaranthus* accessions, we performed Bayesian clustering using STRUCTURE v. 2.3.4. The optimal number of genetic clusters (K) was inferred using the ΔK method described by Evanno et al. (2005). The ΔK plot revealed a pronounced peak at K = 2, indicating the presence of two major genetic groups within the dataset (**Figure 2**.). This suggests a strong primary subdivision among the sampled *Amaranthus* individuals, potentially corresponding to wild versus cultivated types or geographic origin. Although the highest ΔK was observed at K = 2, a secondary peak at K = 4 was also noted, suggesting the possibility of finer substructure. Visual inspection of the bar plots for K = 4 revealed additional differentiation within the major clusters, which may reflect regional adaptation or breeding history. These results are consistent with previous studies on *Amaranthus* genetic diversity, highlighting both broad and subtle population structure (**Figures 2. and 3**).

**Figure 2.**
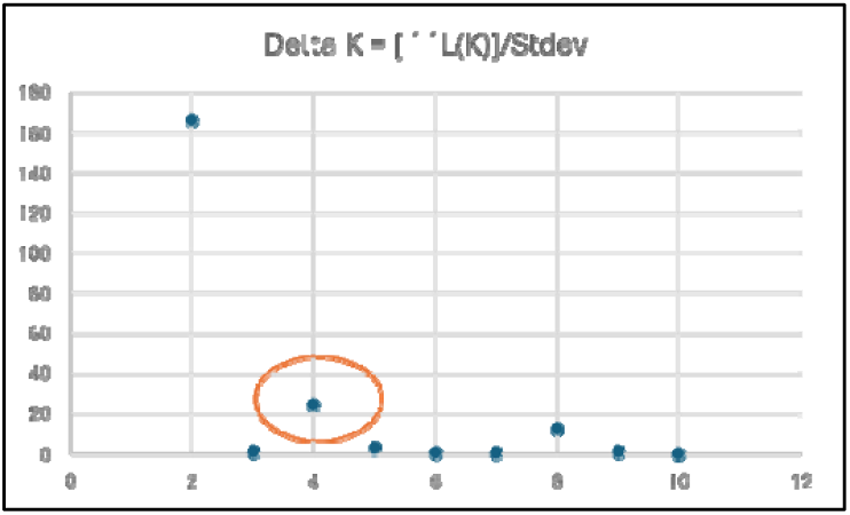

**Figure 3.**
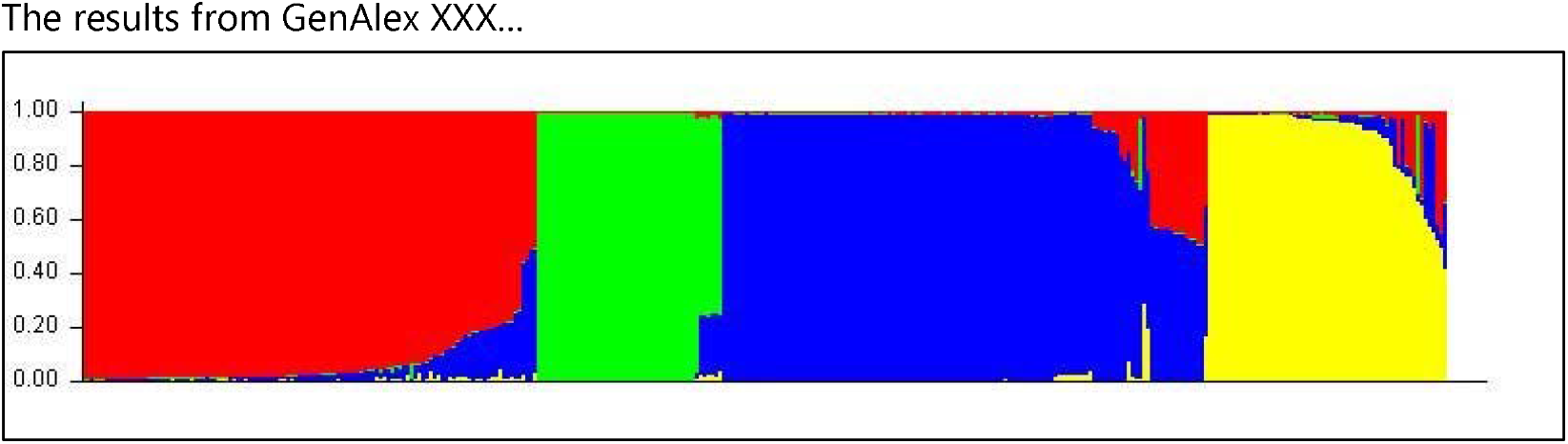

The STRUCTURE bar plot revealed clear patterns of genetic admixture among the *Amaranthus* accessions analyzed (Figures 3. and 4.). Everyone was assigned to one or more genetic clusters, represented by distinct color segments within horizontal bars. The majority of accessions showed strong membership to a single cluster, indicating genetic homogeneity within those groups. However, several samples exhibited mixed ancestry, with multiple color segments suggesting admixture or shared genetic background. Notably, some accessions displayed varying degrees of cluster membership, with some individuals showing near-equal proportions of two or more clusters. These results support the presence of both distinct genetic lineages and admixed individuals within the dataset, consistent with the patterns observed in the ΔK and PCoA analyses. The observed admixture may reflect historical gene flow, hybridization events, or breeding practices that have shaped the genetic landscape of *Amaranthus* populations.

**Figure 4.**
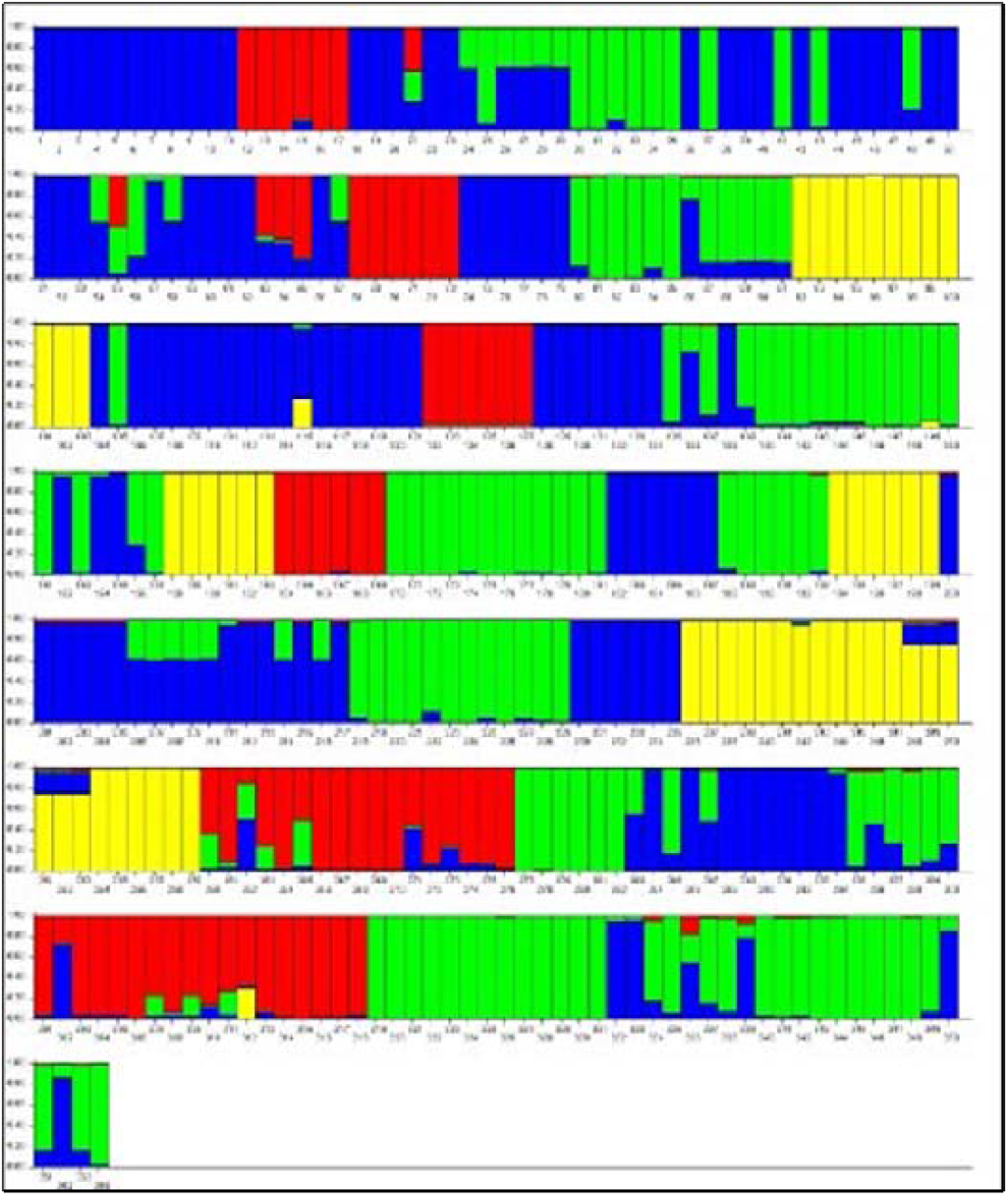

The results from GenAlex XXX…

Microsatellite analysis of *Amaranthus* populations revealed a clear genetic structure among the sampled individuals. STRUCTURE analysis identified four distinct genetic clusters (K = 4), represented by red, green, blue, and yellow in the plot (Figures 3. and 4.). Individuals assigned predominantly to a single cluster displayed high genetic homogeneity, whereas some individuals exhibited mixed ancestry, indicating admixture between clusters. The results suggested that *Amaranthus* populations collected in Burkina Faso structured into four main genetic groups, with varying degrees of admixture. The presence of mixed ancestry individuals indicates ongoing or historical gene flow, while distinct cluster assignments reflect strong genetic differentiation among the major population groups (**Figures 3. and 4**.).

The Principal Coordinates Analysis (PCoA) revealed clear genetic and morphological differentiation among the *Amaranthus* accessions (Figure 5.). Distinct and mixed clusters corresponding to *A. cruentus, A. hypochondriacus*, and *A. dubius* were observed, indicating strong species-level separation. Within these species, morphotype-based groupings emerged, particularly among accessions with corymb or grappe inflorescences and varying leaf shapes. Group characterized by lanceolate leaves and light green morphotypes, formed a cohesive cluster, while accessions with oval-lanceolate leaves and dark green morphotypes showed partial overlap, suggesting intermediate or admixed genotypes. Several accessions, including VBPC_dark green and VSFL Noir, occupied transitional positions between clusters, potentially reflecting hybridization or shared ancestry. These results underscore the utility of PCoA in resolving both taxonomic boundaries and intra-species variation, and they complement the STRUCTURE analysis by highlighting finer-scale diversity shaped by morphological traits and breeding history (**Figure 5.**).

**Figure 5.**
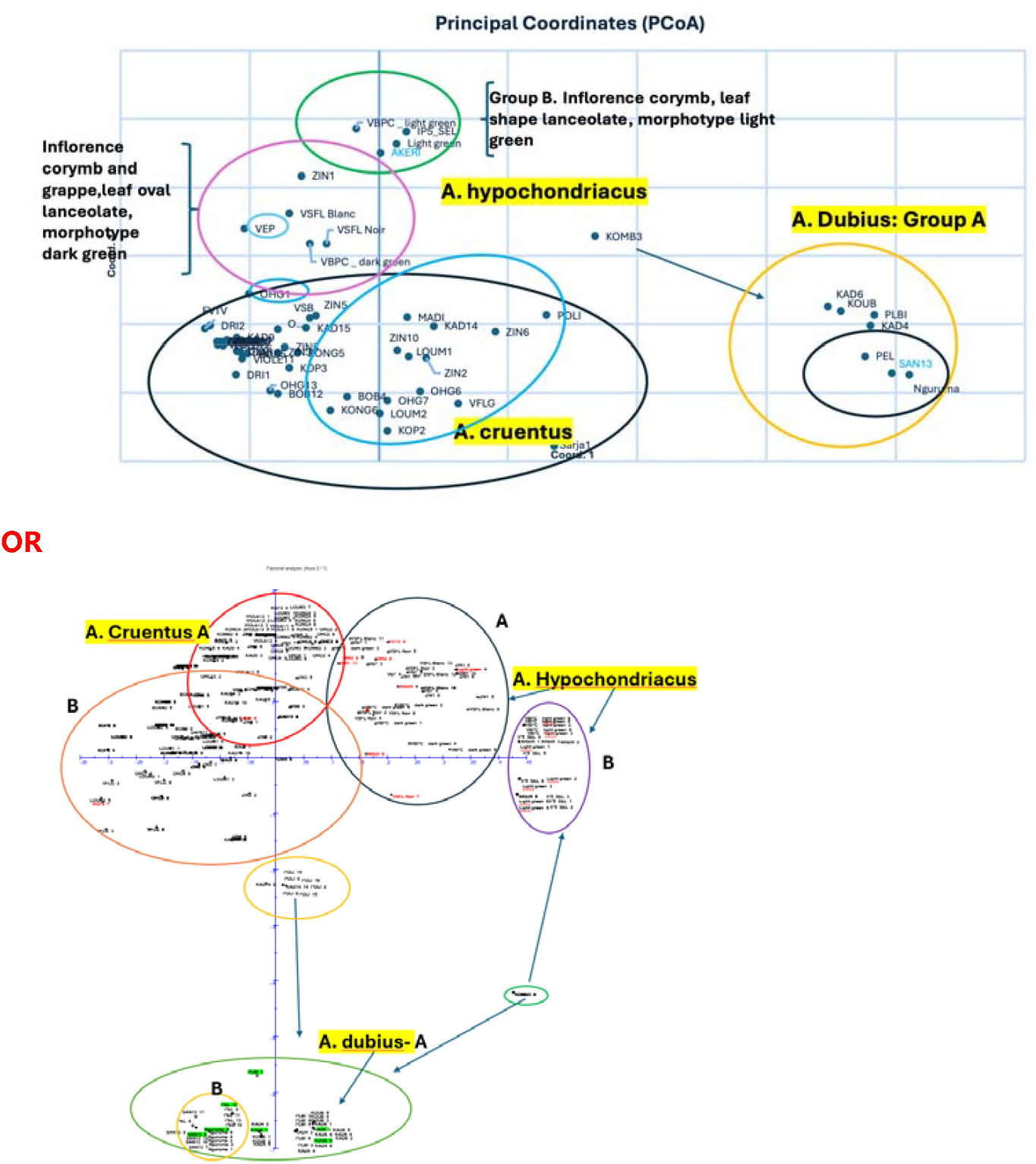
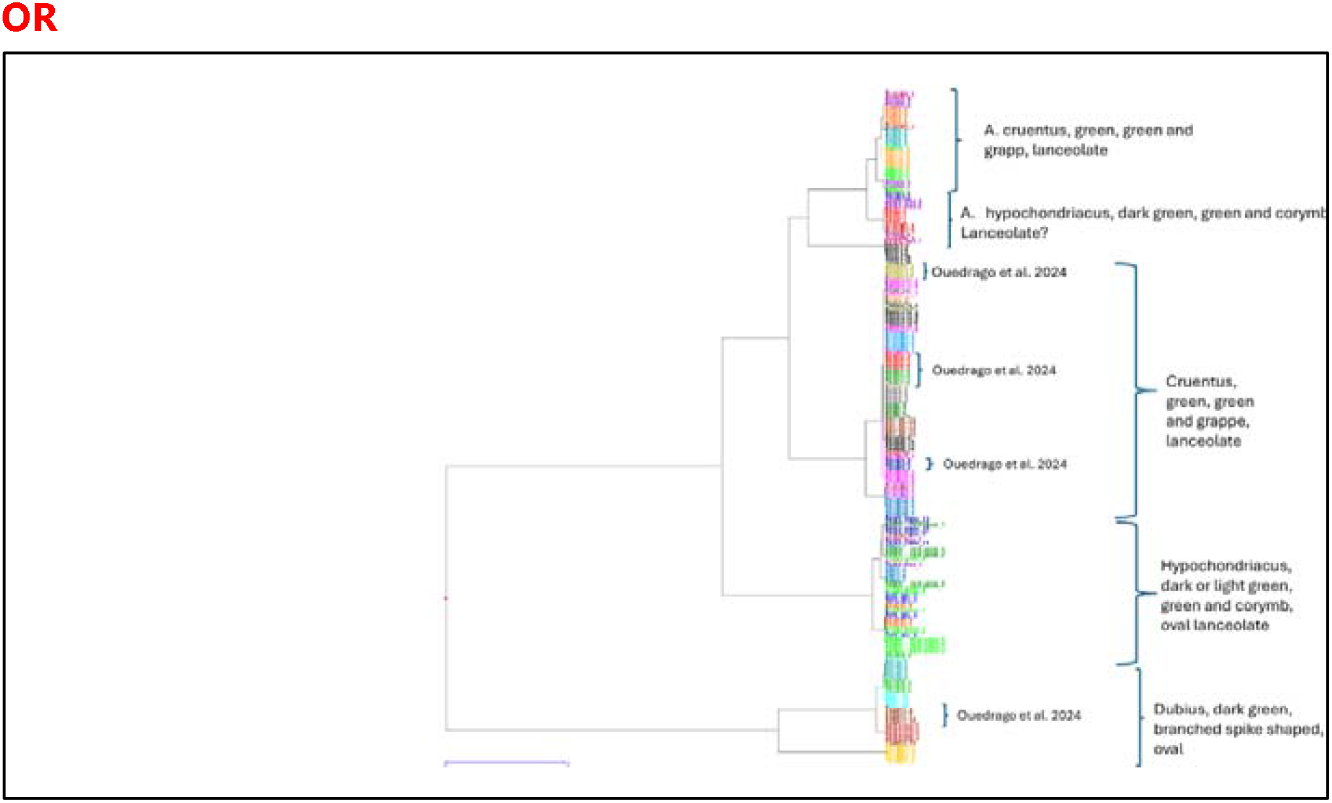

The GBS analysis provided high-resolution insights into the genetic relationships among *Amaranthus* accessions, revealing complex patterns of diversity and structure beyond those detected by SSR markers **(Figures 4. and 5.)**. Hierarchical clustering based on GBS (**Figure 6**.) identified multiple subgroups within species, particularly within *A. cruentus* and *A. hypochondriacus*, where accessions were differentiated by leaf morphology, inflorescence type, and pigmentation. The dendrogram showed clear separation of *A. dubius* accessions, characterized by branched spike-shaped inflorescences and oval leaves, while transitional genotypes exhibited intermediate placement between major clusters, suggesting historical introgression or admixture.

**Figure 6.**
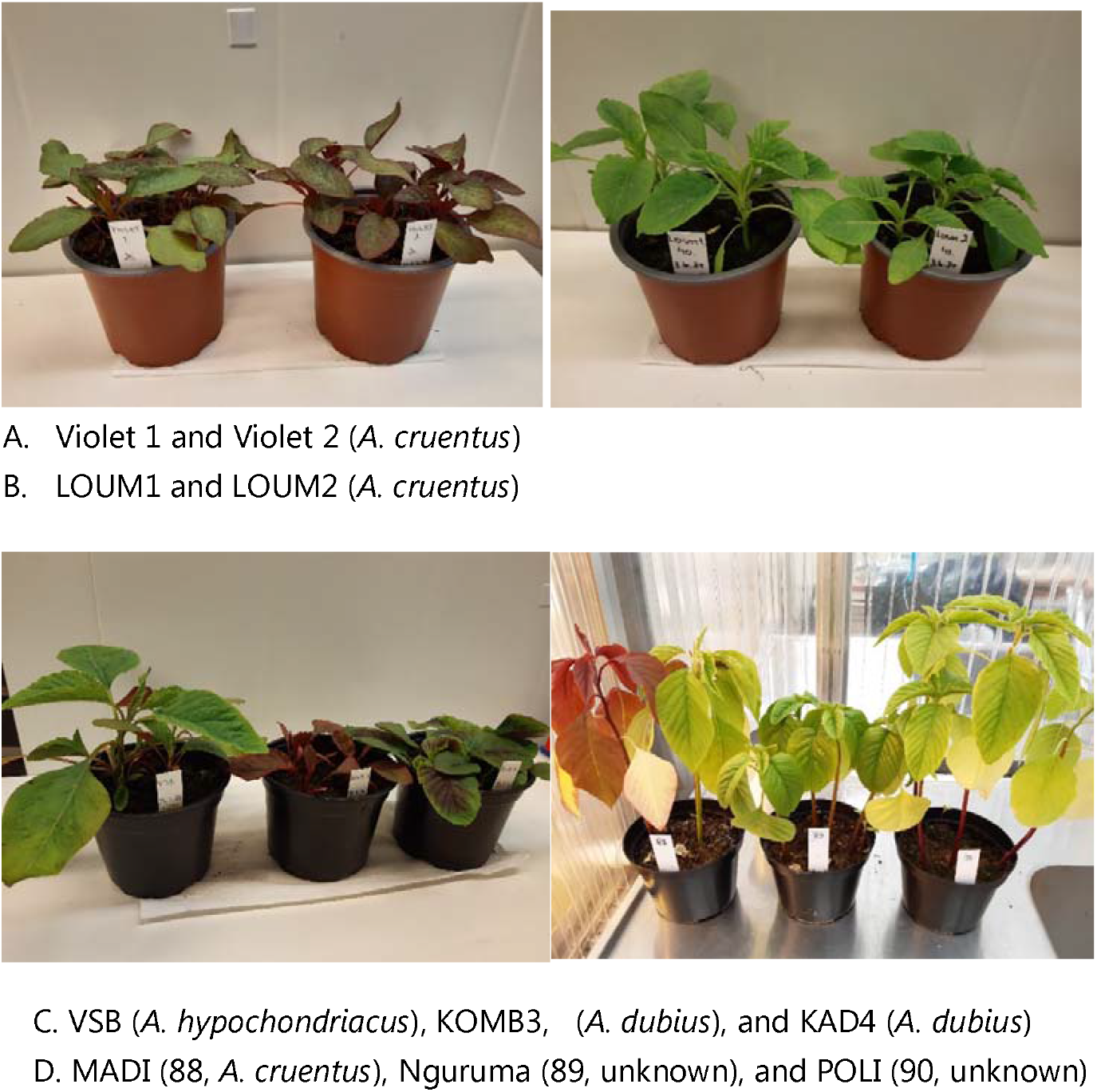
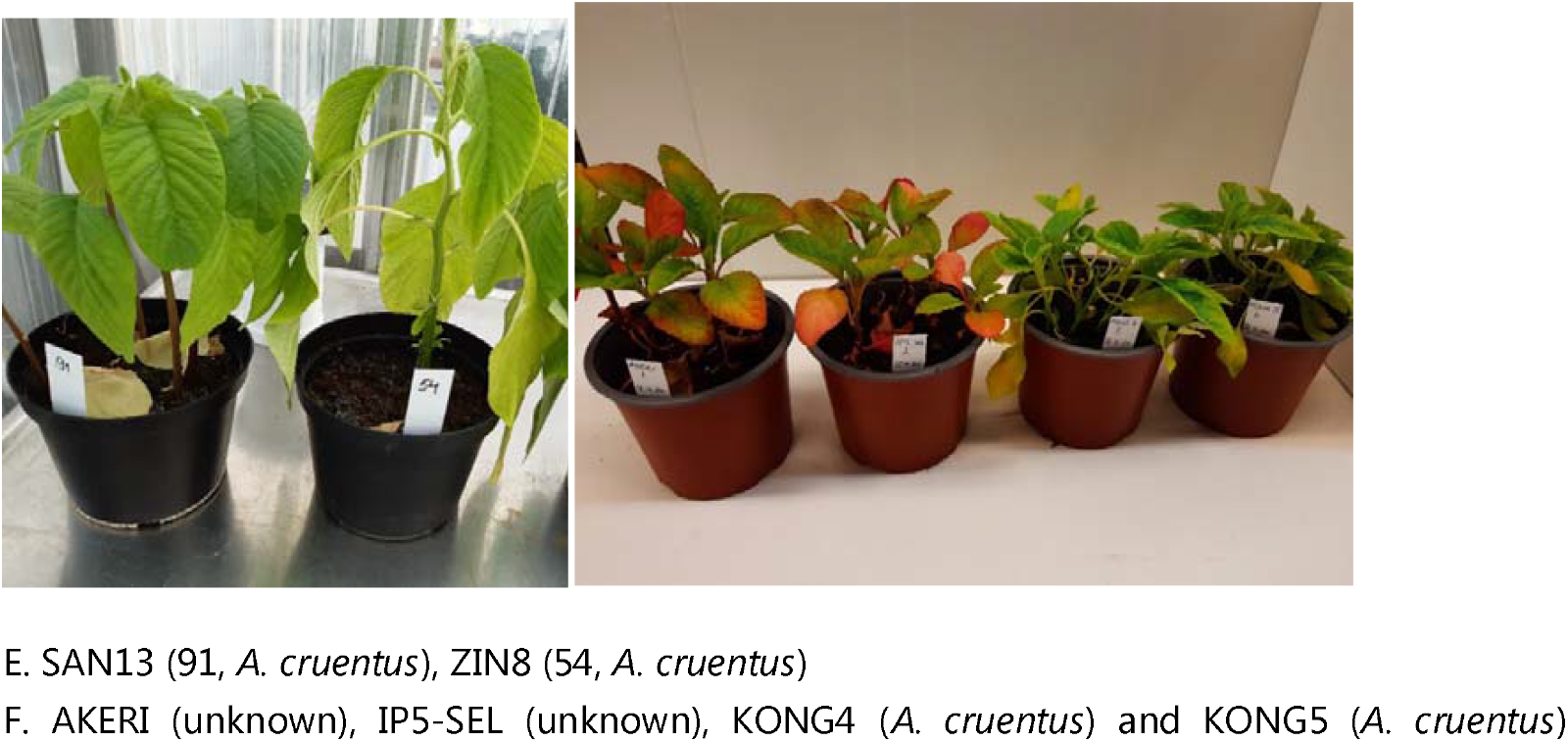

**Figure 6.**
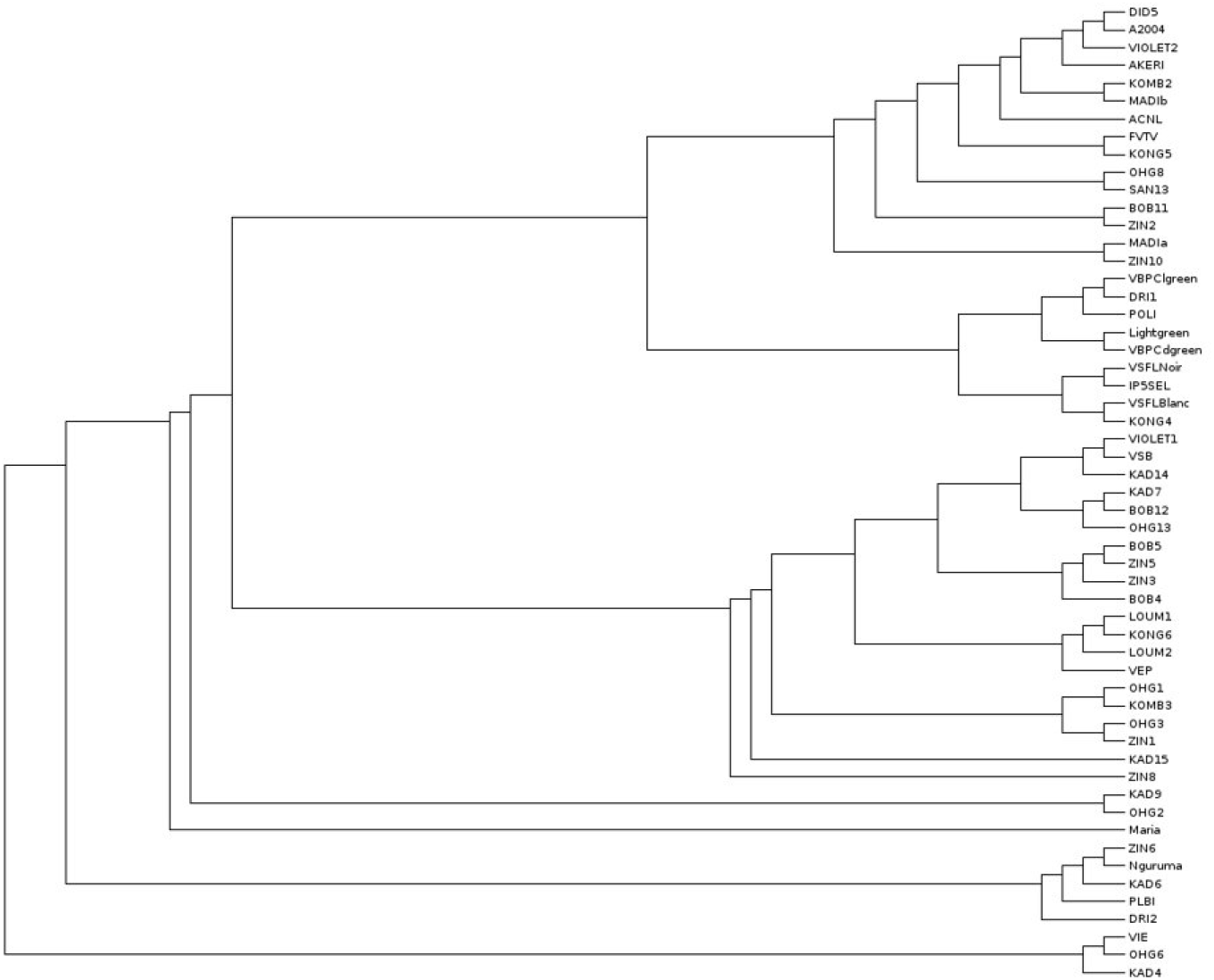
Amaranth tree built from GBS results. Three random samples were used to create a mock reference.

## Discussion

Burkina Faso is a landlocked country in sub-Saharan Africa, and it is one of the least developed countries in the world and suffering from political instability (Rokka et al. 2025). The global food system has been dependent on only a few plant and animal species and strains and there are considerable challenges in achieving food and nutrition security in less-developed countries, such in Burkina Faso. Therefore, also NUS (neglected and underutilized species) have an essential role for maintenance of food security. Amaranth is one of such crops with high genetic diversity and it is also well adapted to drought and marginal lands.

Detailed diversity studies, using molecular markers, genomic sequencing, and phenotypic evaluation, allow breeders to detect unique alleles and gene pools that may carry adaptive traits in marginal crops, such as amaranths. Molecular markers are essential tools in genomics-assisted breeding, facilitating the characterization and utilization of crop genetic resources for efficient varietal improvement (Varshney et al. 2009). Various marker systems, including RFLPs, SSRs, RAPDs, ISSRs, AFLPs, and SNPs, have been utilized in genetic research and breeding, with SNPs being the most abundant and uniformly distributed, providing highly effective and precise genetic analysis (Grover & Sharma 2014; Vignal et al. 2002).

By incorporating underutilized genetic resources into breeding programs, scientists can broaden the genetic base of cultivated varieties and reduce vulnerability to environmental change. In addition, exploring genetic variation among *Amaranthus* contributes to conservation strategies by revealing threatened or genetically unique populations. Maintaining these genetic reservoirs is especially important in the context of climate change, as many wild genotypes possess traits that could be critical for future crop resilience. Overall, genetic diversity studies not only deepen our understanding of the evolutionary dynamics within this genus but also provide the foundation for sustainable crop improvement and long-term food security.

Ouedraogo et al. (2021) has revealed significant phenotypic diversity based on morphological characters in local amaranths of Burkina Faso. Those studies have shown great morphological variability within cultivated *Amaranthus* species, and the authors identified seven morphotypes, when the main classification criteria were the color and shape of leaves and inflorescence (Ouédraogo et al. 2021). Statistical analysis showed that specific traits such as plant height, stem diameter, number of branches, and flowering time were key differentiators between *Amaranthus* species. This diversity among amaranth population shows variations in stem and leaf colors, leaf shape and size, and inflorescence color ranging from maroon to crimson, and the amaranth plants can grow between 1 and 2.5 meters tall in Burkina Faso. Seedlings of amaranths were found in three color variants: green (85%), purple (12.5%), and violet (2.5%). The root system was shallow and pivoting, with observed root colors including red, light pink, and white. Leaf color at the heading stage ranged from green to violet, with light green being the most common. Leaves in most amaranths had a lanceolate shape, while others were oval or lanceolate oval (Ouédraogo et al. 2021).

In the present study, the genetic analysis using SSR and SNP markers by GBS from *Amaranthus* accessions of Burkina Faso revealed complementary insights into their population structure, different species, and their diversity. The analysis of microsatellite variation in the *Amaranthus* dataset revealed a clear genetic subdivision, with STRUCTURE identifying a pronounced ΔK peak at K = 2. This indicated the presence of two major genetic groups, consistent with a primary division that may reflect wild versus cultivated forms or broad geographic differentiation. A secondary signal at K = 4 suggested additional substructure, pointing to finer-scale genetic differentiation within the broader clusters. Such hierarchical structuring is typical of crop species with complex domestication histories, where major divisions are complemented by local adaptation and breeding practices.

Comparable results have been reported by Ouedraogo et al. (2024) in Burkina Faso, who genetically analyzed 72 *Amaranthus* accessions using 11 SSR markers. Their study also identified two primary genetic groups, with further differentiation observed among cultivated accessions. The concordance between our findings and theirs underscores the robustness of SSR markers in capturing both broad and subtle genetic structure in *Amaranthus*. Importantly, the presence of secondary substructure in both datasets highlights the role of regional adaptation, farmer selection, and breeding history in shaping genetic diversity. From a conservation perspective, the detection of two major genetic groups suggests that strategies should aim to preserve representatives from both clusters to maintain overall diversity. The finer substructure observed at K = 4 emphasizes the need to conserve local landraces and region-specific cultivars, which may harbor unique alleles of adaptive significance. For breeding programs, the existence of distinct genetic groups provides opportunities to exploit heterosis by crossing divergent parents, while the recognition of substructure can guide the selection of germplasm adapted to specific environments. Taken together, our results contribute to a growing body of evidence that *Amaranthus* genetic diversity is structured at multiple levels. The agreement with previous studies in Burkina Faso strengthens confidence in these patterns and highlights the importance of integrating molecular data with agronomic and geographic information. Future work combining SSR data with high-throughput genomic approaches will further refine our understanding of *Amaranthus* population structure and enhance its utilization in breeding and conservation.

Genotyping-by-sequencing (GBS) has emerged as a powerful tool for population genetic analysis in *Amaranthus* and other non-model plant taxa because it enables simultaneous discovery and genotyping of thousands of genome-wide single nucleotide polymorphisms (SNPs) at a relatively low cost and high throughput (Elshire et al., 2011; Chauhan et al. 2025). Unlike traditional marker systems that require prior development of species-specific assays. GBS integrates marker discovery and genotyping in a single workflow and facilitates robust SNP identification in species with limited genomic resources (Elshire et al., 2011; Fischer et al., 2025; Peterson et al., 2014). This feature is especially advantageous for *Amaranthus*, a genus characterized by high genetic diversity, frequent hybridization, and complex species relationships, where broad genomic coverage allows fine-scale resolution of population structure, admixture, and phylogenetic relationships (Wu & Blair 2017). High-density SNP datasets generated by GBS improve estimates of genetic differentiation (e.g., F_ST), enhance clustering and principal component analyses, and increase power to detect subtle genetic structuring compared with low-density markers (Sonah et al., 2013; Elshire et al., 2011). Moreover, the scalability of GBS permits efficient genotyping of large numbers of individuals, enabling comprehensive assessments of intra- and inter-population diversity, which is critical for evolutionary studies, germplasm characterization, and breeding applications in *Amaranthus* (Maughan et al., 2017). While bioinformatic challenges and missing data can necessitate careful data processing, the flexibility and genome-wide scope of GBS make it particularly well-suited for population genetic analyses in diverse and under-studied plant genera.

The GBS analysis in the present study provided high-resolution insights into the genetic relationships among *Amaranthus* accessions, revealing complex patterns of diversity and structure beyond those detected by SSR markers. Hierarchical clustering based on genome-wide SNP data identified multiple subgroups within species, particularly within *A. cruentus* and *A. hypochondriacus*, where accessions were differentiated by leaf morphology, inflorescence type, and pigmentation. The dendrogram showed clear separation of *A. dubius* accessions, characterized by branched spike-shaped inflorescences and oval leaves, while transitional genotypes exhibited intermediate placement between major clusters, suggesting historical introgression or admixture. Our results highlight the power of GBS to resolve fine-scale genetic structure and uncover subtle differentiation linked to morphological traits and breeding history. The genome-wide coverage of GBS may have the power to identify QTLs associated with adaptation, making it a valuable tool for germplasm characterization and crop improvement in *Amaranthus species*.

Amaranth is one of the underutilized plant species with strong potential to help prevent famine in developing countries. This study also revealed that the Sudanian and Sahelian regions still harbor a wealth of underexplored plant genetic resources. Due to its high nutritional value, including notable protein and fiber content, amaranth may also appeal to developed countries as a novel protein source for Western populations.

## Acknowledgements

Marja-Riitta Arajärvi, Sirpa Moisander and Anneli Virta are thanked for their technical support in Luke Genomics laboratory, Jokioinen, Finland. All the greenhouse staff are appreciated for taking care of Amaranthus plants. The study was funded by DeSIRA, a European Union initiative supporting agriculture and food systems in low and middle-income countries.

